# HyperGen: Compact and Efficient Genome Sketching using Hyperdimensional Vectors

**DOI:** 10.1101/2024.03.05.583605

**Authors:** Weihong Xu, Po-Kai Hsu, Niema Moshiri, Shimeng Yu, Tajana Rosing

**Affiliations:** Department of Computer Science and Engineering, University of California San Diego, La Jolla, CA 92093, USA; School of Electrical and Computer Engineering, Georgia Institute of Technology, Atlanta, GA 30332, USA

## Abstract

**Motivation:** Genomic distance estimation is a critical workload since exact computation for whole-genome similarity metrics such as Average Nucleotide Identity (ANI) incurs prohibitive runtime overhead. Genome sketching is a fast and memory-efficient solution to estimate ANI similarity by distilling representative *k*-mers from the original sequences. In this work, we present HyperGen that improves accuracy, runtime performance, and memory efficiency for large-scale ANI estimation. Unlike existing genome sketching algorithms that convert large genome files into discrete *k*-mer hashes, HyperGen leverages the emerging hyperdimensional computing (HDC) to encode genomes into quasi-orthogonal vectors (Hypervector, HV) in high-dimensional space. HV is compact and can preserve more information, allowing for accurate ANI estimation while reducing required sketch sizes. In particular, the HV sketch representation in HyperGen allows efficient ANI estimation using vector multiplication, which naturally benefits from highly optimized general matrix multiply (GEMM) routines. As a result, HyperGen enables the efficient sketching and ANI estimation for massive genome collections.

**Results:** We evaluate HyperGen’s sketching and database search performance using several genome datasets at various scales. HyperGen is able to achieve comparable or superior ANI estimation error and linearity compared to other sketch-based counterparts. The measurement results show that HyperGen is one of the fastest tools for both genome sketching and database search. Meanwhile, HyperGen produces memory-efficient sketch files while ensuring high ANI estimation accuracy.

**Availability:** A Rust implementation of HyperGen is freely available under the MIT license as an open-source software project at https://github.com/wh-xu/Hyper-Gen. The scripts to reproduce the experimental results can be accessed at https://github.com/wh-xu/experiment-hyper-gen.

## 1 Introduction

In recent years, the burgeoning field of genomics has been revolutionized by the advent of high-throughput sequencing technologies (Soon *et al*., 2013), leading to exponential growth in genomic data (Stephens *et al*., 2015). This deluge of data presents a significant challenge for traditional genomic analysis methods, particularly in terms of computational efficiency and storage requirements. Calculating the Average Nucleotide Identity (ANI) similarity of genome files is the key step for various downstream workloads in genome analysis, such as large-scale database search (Chaumeil *et al*., 2022), clustering (Parks *et al*., 2020), and taxonomy analysis (Hernández-Salmerón *et al*., 2023). Traditional BLAST-based methods (Kurtz *et al*., 2004; Lee *et al*., 2016) rely on base-level alignment to perform accurate ANI calculations. However, the alignment process is computationally expensive and requires hours or days to calculate ANIs. The slow speed of alignment-based approaches has become a major bottleneck for large-scale genome analysis.

Several state-of-the-art works have tried to speed up large-scale genome analysis by approximating the genome similarity using more efficient data structures. These works can be categorized into two types: mapping-based and sketch-based approaches as follows. FastANI (Jain *et al*., 2018) and Skani (Shaw and Yu, 2023) are two representative mapping-based algorithms that leverage *k*-mer-based alignment for ANI estimation. FastANI is built upon the Mashmap sequence mapping algorithm (Jain *et al*., 2017) and achieves a significant speedup compared to the alignment-based baseline (Kurtz *et al*., 2004). Skani uses the sparse chaining to increase the sensitivity of the mapping, further improving accuracy and efficiency of ANI estimation. However, both FastANI and Skani suffer from high memory consumption. For example, Skani needs to store indexing files with a storage size comparable to the original dataset. FastANI encounters out-of-memory issues on large datasets as reported in (Shaw and Yu, 2023).

In this work, we focus on the “*genome sketching*,” which is regarded as a promising solution to address the aforementioned challenges because it significantly reduces storage size while providing satisfactory accuracy of estimation (Hernández-Salmerón *et al*., 2023). Unlike alignment-based or mapping-based tools (Kurtz *et al*., 2004; Lee *et al*., 2016; Jain *et al*., 2018; Shaw and Yu, 2023) that require expensive computation or large memory space, sketch-based approaches (Ondov *et al*., 2016; Brown and Irber, 2016; Baker and Langmead, 2019, 2023) only preserve the most essential features of the genome (called the “sketch”). The sketch’s compact representation enables rapid and efficient ANI approximation for genome files. Mash (Ondov *et al*., 2016) and Sourmash (Brown and Irber, 2016) represent groundbreaking efforts to use MinHash (Broder, 1997) and FracMinHash (Hera *et al*., 2023; Irber *et al*., 2022) to estimate genomic similarity, respectively. Bindash (Zhao, 2019) improves the accuracy of ANI estimation over Mash by adopting the one-permutation rolling MinHash with optimal densification (Shrivastava, 2017). Dashing 2 (Baker and Langmead, 2023) utilizes the SetSketch data structure (Ertl, 2021) and incorporates multiplicities to produce memory-efficient genome sketches and accurate estimation of ANI.

### 1.1 Motivation

By transforming raw genome data into more compact data structures, genome sketching represents a paradigm shift in bioinformatics, paving the way for more scalable and rapid genomic analyses in the era of big data. Recent studies on hyperdimensional computing (HDC) have demonstrated the effectiveness of using HDC to accelerate bioinformatics workloads, such as pattern matching (Zou *et al*., 2022; Kang *et al*., 2023; Kim *et al*., 2020; Shahroodi *et al*., 2022) and spectral clustering (Xu *et al*., 2023).

#### 1.1.1 Limitations of existing HDC/SimHash-related search algorithms

Table 1 summarizes the key features of state-of-the-art tools that utilize HDC or SimHash algorithms. GenieHD (Kim *et al*., 2020), BioHD (Zou *et al*., 2022), and Demeter (Shahroodi *et al*., 2022) are three representative HDC-based tools. Due to the limitation of *N* -gram binding-based encoding, existing HDC tools for genome search only supports short genomes sequences with length ≤ 200. However, they require very large sketch HV dimension (10k to 100k) to achieve good accuracy, which degradates the overall efficiency. The *N* -gram binding-based encoding shows high computational complexity. In comparison, HyperGen adopts a more efficiency encoding method that combines FracMinHash and HDC aggregation.

**Table 1.** Comparison for related works for genome search and seed matching.

| Algorithm / Tool | GenieHD<br>(Kim et al., 2020) | BioHD<br>(Zou et al., 2022) | Demeter<br>(Shahroodi et al., 2022) | BLEND<br>(Firtina et al., 2023) | HyperGen<br>(This work) |
| --- | --- | --- | --- | --- | --- |
| Encoding method | $N$-gram HDC binding | $N$-gram HDC binding | $N$-gram HDC binding | SimHash | FracMinHash + HDC |
| Supported sequence length | $\leq 200$ | $\leq 200$ | $\approx 150$ | 150-20k | Arbitrary |
| Sketch dimension | 100k | 10k-40k | 40k | 30-50* | 2k-8k |
| Support ANI estimation? | No | No | No | No | Yes |
| Supported application | Containment search | Containment search | Containment search | Seed matching | ANI-based search and clustering |

\* Sketch dimension for each seed.

Meanwhile, existing HDC-based tools do not support ANI estimation and ANI-based search. They can only check the containment of given query. These drawbacks limit their downstream applications. The other related work is BLEND (Firtina *et al*., 2023) that uses SimHash to encode genome seeds. The difference includes: 1. HyperGen and BLEND are for different tasks. BLEND is used for seed matching while HyperGen is for more general-purpose ANI estimation and database search, 2. Compared to HyperGen, BLEND uses much smaller sketch dimension for each seed.

#### 1.1.2 Opportunities and Limitations of DotHash

Recent DotHash (Nunes *et al*., 2023) shows superior space and computational efficiency for the Jaccard similarity estimation. DotHash leverages the HDC-based random indexing (Sahlgren, 2005; Kanerva *et al*., 2000) and is originally designed for fast set intersection estimation. The main difference between DotHash and MinHash lies in the format of generated sketch: MinHash represents a sketch as a hash set with discrete values, while DotHash represents a sketch with a nonbinary vector of high dimension. DotHash’s vector representation of the sketch achieves faster processing speed since it can fully exploit the low-level hardware parallelism (such as CPU’s Single Instruction Multiple Data (SIMD) and GPU) optimized for vector processing.

However, DotHash still suffers from two major limitations that hinder its application to genome sketching. First, DotHash is only applicable to non-genome data since it lacks an effective *k*-mer sampling strategy to generate genomic sketches. Second, DotHash uses high-precision floating point numbers to represent random vectors, exhibiting large runtime overhead and slow speed. Our goal in this work is using HDC (Kanerva, 2009; Nunes *et al*., 2023) to achieve better tradeoffs between ANI estimation accuracy, runtime performance, and memory efficiency over previous sketch-based tools (Ondov *et al*., 2016; Brown and Irber, 2016; Baker and Langmead, 2023).

### 1.2 Contributions

In this work, we propose HyperGen, a novel tool for efficient genome sketching and ANI estimation. HyperGen exploits the emerging HDC (similar to DotHash (Nunes *et al*., 2023)) to boost genomic ANI calculation. Specifically, we optimize DotHash’s efficiency by converting the sketch generation process into a low bit-width integer domain. This allows us to represent the genome sketch using the high-dimensional vector (HV) at the cost of negligible runtime overhead. Based on the HV sketch, we propose an approach to estimate the Jaccard similarity using vector matrix multiplication. We also introduce a lossless compression scheme using bit-packing to further reduce the sketch size.

We benchmark HyperGen against several state-of-the-art tools (Jain *et al*., 2018; Ondov *et al*., 2016; Baker and Langmead, 2023; Kurtz *et al*., 2004). For ANI estimation, HyperGen demonstrates comparable or lower ANI estimation errors compared to other baselines across different datasets.

For generated sketch size, HyperGen achieves 1.8× to 2.7× sketch size reduction as compared to Mash (Ondov *et al*., 2016) and Dashing 2 (Baker and Langmead, 2023), respectively. HyperGen also enjoys the benefits of the modern hardware architecture optimized for vector processing. HyperGen shows about 1.7× sketch generation speedup over Mash and up to 4.3× search speedup over Dashing 2. To the best of our knowledge, HyperGen offers the optimal trade-off between speed, accuracy, and memory efficiency for ANI estimation.

## 2 Methods

### 2.1 Preliminaries

Fast computation of Average Nucleotide Identity (ANI) is pivotal in genomic data analysis (microbial genomics to delineate species), as ANI serves as a standardized and genome-wide measure of similarity that helps facilitate genomic data analysis. Popular approaches to calculate ANI include: alignment (Kurtz *et al*., 2004; Lee *et al*., 2016), mapping (Jain *et al*., 2018; Shaw and Yu, 2023), and sketch (Ondov *et al*., 2016; Brown and Irber, 2016; Baker and Langmead, 2019, Baker and Langmead 2023). However, base-level alignment-based and *k*-mer-level mapping-based methods involve either time-consuming pairwise alignments or memory-intensive mappings. In the following sections, we focus on the sketch-based ANI estimation with significantly better efficiency.

#### 2.1.1 MinHash and Jaccard Similarity

Existing sketh-based approaches (Ondov *et al*., 2016; Brown and Irber, 2016; Baker and Langmead, 2019, 2023) do not directly compute ANI. Instead, they compute the Jaccard similarity (Ondov *et al*., 2016), which is used to measure the similarity of two given *k*-mer sets. Then the Jaccard similarity is converted to ANI as shown in Eq. (8). The conversion between Jaccard similarity and ANI is computationally trivial, so most efforts in previous works (Ondov *et al*., 2016; Brown and Irber, 2016; Baker and Langmead, 2019, 2023) are to find more efficient and accurate ways to estimate Jaccard similarity.

Without loss of generality, we denote *k*-mer as consecutive substrings with length *k* of the nucleotide alphabet, *e*.*g*. ∑^*k*^ = {*A, G, C, T*}^*k*^. 𝒮_*k*_(*X*) denotes the set of *k*-mers sampled from genome sequence *X* based on a given condition. HyperGen uses *k*-mer’s hash to represent 𝒮_*k*_(*X*) for better efficiency. Therefore, the Jaccard similarity for two sequences, *A* and *B*, can be computed as follows:

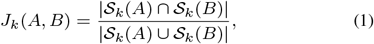

where *J*_*k*_(*A, B*) ∈ [0, 1] is the Jaccard similarity indicating the overlap between *k*-mer sets of two sequences. Note that HyperGen uses canonical *k*-mers by default.

A straightforward idea to sample *k*-mer sets in Eq. (1) is to keep all

*k*-mers. However, this incurs prohibitive complexity since all unique *k*-mers need to be stored. The resulting complexity is 𝒪 (*L*) for a sequence of length *L*. To alleviate the complexity issue, Mash (Ondov *et al*., 2016) and its variants (Liu and Koslicki, 2022; Jain *et al*., 2017) use MinHash (Broder, 1997) to approximate the Jaccard similarity by only preserving a tiny subset of *k*-mers. In particular, Mash keeps *N k*-mers that have the smallest hash values *h*(·). In this case, the Jaccard similarity is estimated as:

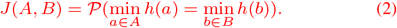

Here, using MinHash helps to reduce the sketch complexity from 𝒪 (*L*) to a constant 𝒪 (*N*). The sampled *k*-mer set 𝒮_*k*_(*X*) that stores *N* smallest *k*-mer hash values is regarded as the genome file sketch required for ANI estimation.

#### 2.1.2 Jaccard Similarity using DotHash

A recent work (Nunes *et al*., 2023) demonstrates that the speed and memory efficiency of Jaccard similarity approximation can be improved by using the DotHash based on Random Indexing (Sahlgren, 2005). The key step to compute Jaccard similarity in Eq. (1) is computing the cardinality of set intersection |*A* ∩ *B*| while the cardinality of set union can be calculated through |*A* ∪ *B*| = |*A*| + |*B*| − |*A* ∩ *B*|.

In DotHash, each element of the set is mapped to a unique *D*-dimensional vector in real number using the mapping function *ϕ*(*x*). Each set is expressed as an aggregation vector a ∈ ℝ^*D*^ such that

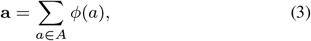

where the aggregation vector sums all the elements’ vectors generated by the mapping function *ϕ*(*x*). One necessary constraint for function *ϕ*(*x*) is: the generated vectors should satisfy the quasi-orthogonal properties:

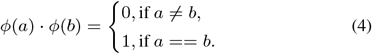

The quasi-orthogonal property in Eq. (4) can be visualized in Supplementary Fig. 1. DotHash (Nunes *et al*., 2023) uses a pseudo random number generator (RNG) as the mapping function *ϕ*(*x*) because the RNG can generate uniform and quasi-orthogonal vectors in an efficient manner.

Using the quasi-orthogonal properties, the cardinality approximation for set intersection is transformed into the dot product of two aggregation vectors:

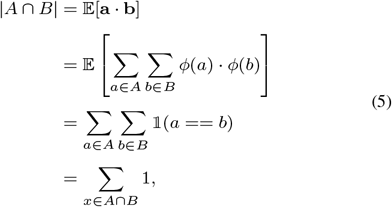

where those vectors not in the set intersection (*a* ≠ *b*) have no contribution to the inner product due to their quasi-orthogonality as in Eq. (4). DotHash effectively aggregates all elements in a set to form an aggregation vector with *D* dimension. The space and computational complexity of set cardinality estimation is 𝒪 (*D*). Moreover, the computation process of DotHash is highly vectorized and can be easily boosted by existing hardware architecture optimized for general matrix multiply (GEMM).

### 2.2 Proposed HyperGen Sketching

The aforementioned DotHash provides both good accuracy and runtime performance (Nunes *et al*., 2023). However, we observe two major limitations of DotHash: 1. Although DotHash can be used to calculate the cardinality of set intersection, it cannot be applied to genomic sketching because DotHash lacks a *k*-mer sampling module that identifies the useful *k*-mers; 2. The computation and space efficiency can be further optimized because the previous DotHash manages and processes all vectors in floating-point (FP) numbers. The mapping function *ϕ*(*x*) incurs significantly overhead.

We present HyperGen for genomic sketching applications that addresses the limitations of DotHash. Fig. 1 shows the algorithmic overview for (a) Mash-like sketching and (b) HyperGen sketching schemes. The first step of HyperGen is similar to Mash, where both Mash and HyperGen extract *k*-mers by sliding a window through given genome sequences. The extracted *k*-mers are uniformly hashed into the corresponding numerical values by a hash function *h*(*x*). To ensure low memory complexity, most *k*-mer hashes are filtered and only a small portion of them are preserved in the *k*-mer hash set to work as the sketch (or signature) of the associative genome sequence. The key difference is that HyperGen adds a key step, called *Hyperdimensional Encoding for k-mer Hash*, to convert *k*-mer hash values into binary hypervectors (HVs) and aggregate to form the *D*-dimensional sketch HV. To distinguish itself from DotHash, the random vector in HyperGen is named HV. Algorithm 1 summarizes the flow of generating sketch hypervector in HyperGen. In the following sections, we explain the details of HyperGen.

**Fig. 1:**
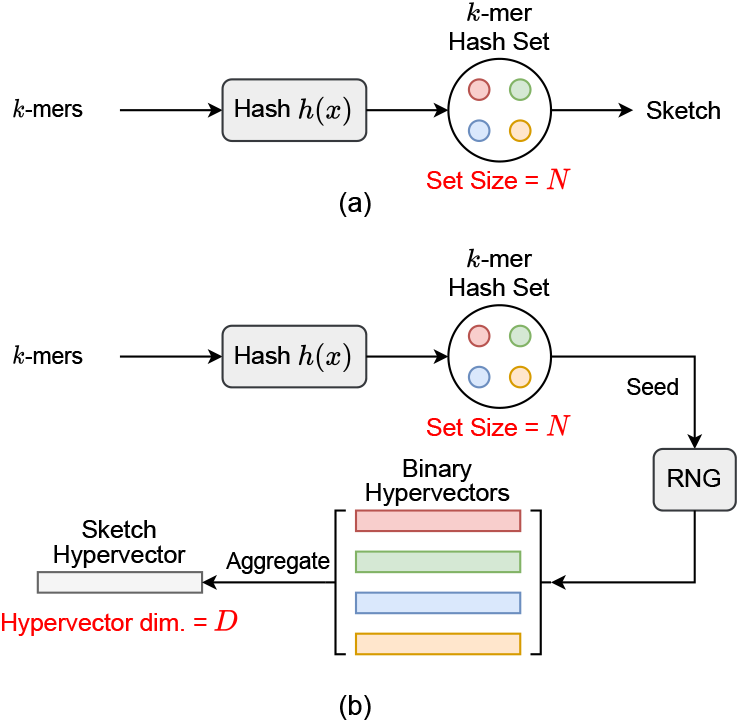
Algorithmic overview for (a) Mash-like sketching, and (b) HyperGen sketching for genome sequences. Mash stores the genome sketch in a *k*-mer hash set with 𝒪 (*N*) complexity while HyperGen aggregates *N k*-mer hashes into a *D*-dimensional sketch HV with 𝒪 (*D*) complexity.

#### 2.2.1 Step 1: *k*-mer Hashing and Sampling

Mash uses MinHash that keeps the smallest *N* hash values as the genome sketch. In comparison, HyperGen adopts a different *k*-mer hashing and sampling scheme. Specifically, HyperGen performs a sparse *k*-mer sampling using FracMinHash (Hera *et al*., 2023; Irber *et al*., 2022) (instead of MinHash in Mash). Given a hash function *h* : ∑^*k*^ → [0, *M*] that maps *k*-mers into the corresponding nonnegative integer, the sampled *k*-mer hash set is expressed as Line 2-4 in Algorithm 1:

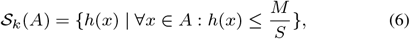

where *M* is the maximum hash value while *S* denotes the scaled factor that determines the density of sampled *k*-mers in the set. FracMinHash has been widely adopted in other tools, such as Sourmash (Brown and Irber, 2016) and Skani (Shaw and Yu, 2023), due to its excellent performance. The advantage of using FracMinHash over MinHash (Broder, 1997) is that it ensures an unbiased estimation of the Jaccard similarity of *k*-mer sets with very dissimilar sizes (Hera *et al*., 2023), providing better approximation quality than MinHash and its variants (Ondov *et al*., 2016; Jain *et al*., 2017). However, FracMinHash usually produces a larger hash set compared to Mash (Hera *et al*., 2023), requiring more memory space. Step 2 in HyperGen alleviates the increased memory issue.

##### Algorithm 1

Generation of sketch hypervector in HyperGen

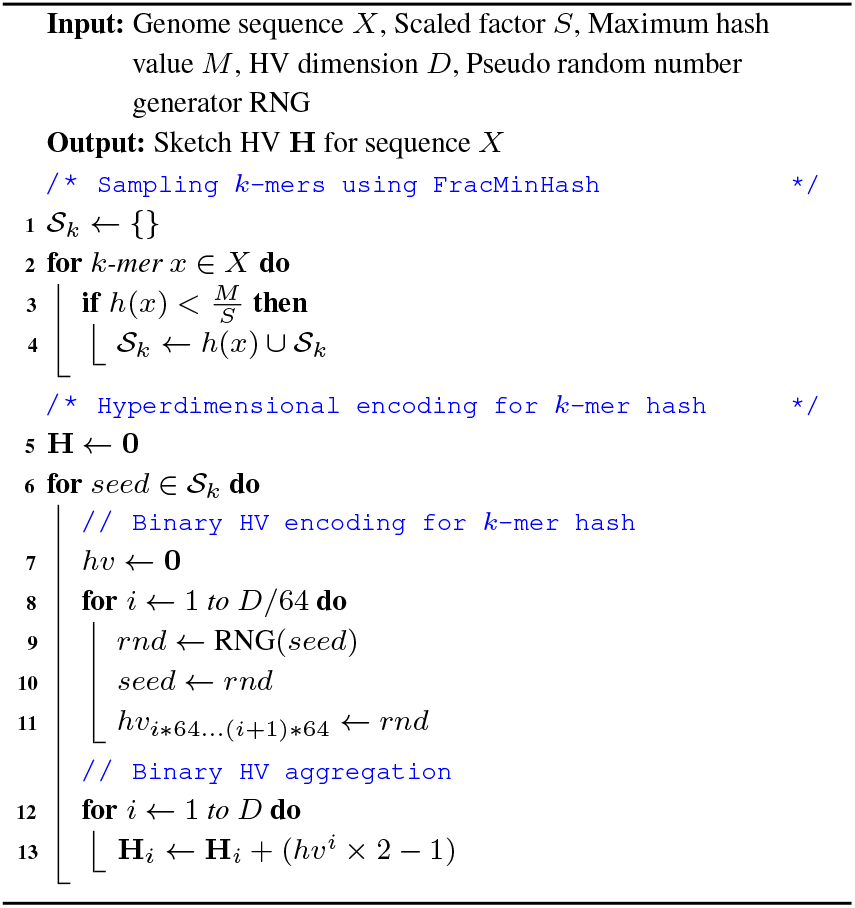

#### 2.2.2 Step 2: Hyperdimensional Encoding for *k*-mer Hash

In Fig. 1-(a), after the *k*-mer hashing and sampling process, Mash-like sketching algorithms (such as Mash (Ondov *et al*., 2016), Sourmash (Brown and Irber, 2016), and Mash Screen (Ondov *et al*., 2019)) directly use the sampled *k*-mer hash set as the sketch to compute the Jaccard similarity for given sequences.

In Fig. 1-(b), HyperGen adds an additional step, called *Hyperdimensional Encoding for k-mer Hash* (Line 5-13 in Algorithm 1), before the sketch is generated. This step essentially converts the discrete and numerical hashes in the *k*-mer hash set to a *D*-dimensional and nonbinary vector, called *sketch hypervector*. The hypervector dimension *D* is normally large (1024 to 8192) to ensure good accuracy. In particular, each hash value in the *k*-mer hash set is uniquely mapped to the associated binary HV *hv* as Line 6-11 of Algorithm 1. HyperGen relied on recursive random bit generation to produce binary HVs of arbitrary length: the *k*-mer hash value is set as the initial seed of the pseudo RNG(*seed*) → *rnd* function. For each iterative step, a 64b random integer *rnd* is generated using *seed*. The generated integer *rnd* is not only assigned to the corresponding bits in *hv*, but is also set as the next *seed*.

The hash function RNG(·) that maps the *k*-mer hash value to the binary HV *hv* is the key component of HyperGen because it determines the speed and quality of genome sketch generation. The following factors should be considered when selecting a good RNG(·) function: 1. The function needs to be fast enough to reduce the additional overhead for sketch generation. 2. The generated random binary HVs need to be able to provide enough randomness (*i*.*e*., the binary HVs are as orthogonal as possible). This is because binary HVs are essentially random binary bit streams that need to be nearly orthogonal to each other to satisfy the quasi-orthogonal requirements. 3. The sketches results should be reproducible (*i*.*e*., the identical bit streams can be generated using the same seed). We adopt a fast and high-quality pseudo RNG^1^ in Rust language (Matsakis and Klock, 2014), which passes two randomness tests: TestU01 and Practrand (Sleem and Couturier, 2020). In this case, we can use the pseudo RNG to stably generate high-quality and reproducible binary HVs.

Fig. 2 shows an example of generating the sketch HVs with dimension *D* = 8 for two genome sequences based on *k*-mer size *k* = 3 and *k*-mer hash set size *N* = 4. Each sampled *k*-mer hash value in the hash set is converted to the corresponding binary HV *hv* ∈ {0, 1}^*D*^ using the function RNG(*x*). Then, all *N* binary HVs are aggregated into a single sketch HV H ∈ ℤ^*D*^ based on the following point-wise vector addition:

**Fig. 2:**
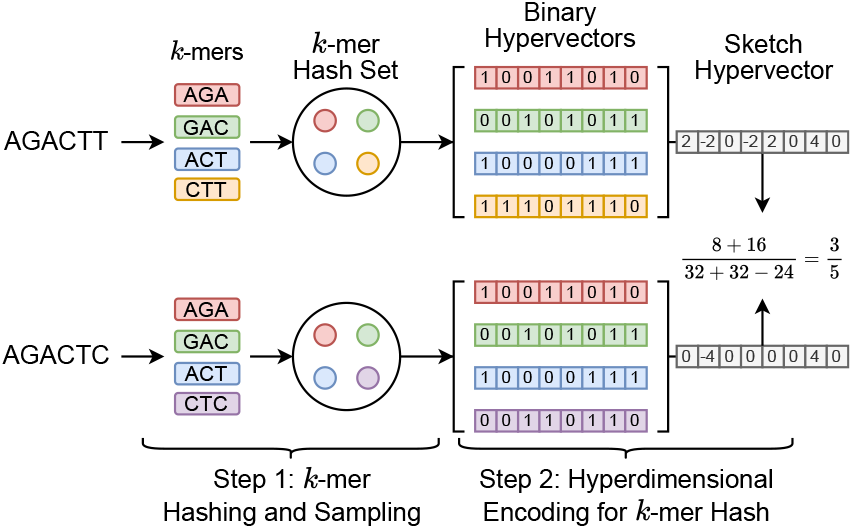
Sketch hypervector generation and set intersection computation in HyperGen. Each *k*-mer with size *k* = 3 first passes through a hash function *h*(*x*). The *k*-mers (*A* = *AGACTT* and *B* = *AGACTC*) are hashed to hash set. Then each *k*-mer hash value is converted into the associated orthogonal binary HV. The set intersection between two *k*-mer hash sets is computed using Eq. (11).

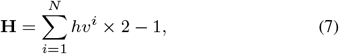

where the binary HV *hv* ∈ {0, 1}^*D*^ is first converted to {−1, +1}^*D*^. *hv*^*i*^ denotes the *i*-th binary HV in the set. Then all binary HVs in the set are aggregated together to create the corresponding sketch HV. Compared to Mash-liked sketching approaches (Ondov *et al*., 2016; Hera *et al*., 2023; Brown and Irber, 2016), HyperGen is more memory efficient because the sketch HV format is more compact with 𝒪 (*D*) space complexity, which is independent of the *k*-mer hash set size *N*. Meanwhile, HyperGen’s hyperdimensional encoding step helps to achieve better ANI similarity estimation quality (see Section 3).

#### 2.2.3 Step 3: ANI Estimation using Sketch Hypervector

The generated sketch hypervector can be used to efficiently estimate the ANI similarity. HyperGen estimates ANI value using the same approach in (Ondov *et al*., 2016). The ANI under the Poisson distribution is estimated as:

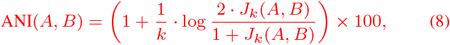

where *J*_*k*_(*A, B*) denotes the Jaccard similarity between genome sequence *A* and sequence *B* while *k* is the *k*-mer size.

Therefore, ANI estimation in HyperGen becomes calculating Jaccard similarity based on sketch HVs. Eq. (1) shows that the intersection size and the set size of two *k*-mer hash sets are the keys to calculating the Jaccard similarity. For *hv*^*i*^ ∈ {−1, +1}^*D*^, the cardinality of a set 𝒮_*k*_(*A*) is computed as follows:

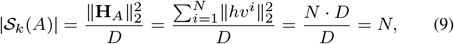

which shows the set cardinality can be computed based on the *L*^2^ norm of sketch HV. The computation of set intersection in HyperGen is similar to DotHash (Nunes *et al*., 2023)’s Eq. (5) because HVs in HyperGen share the same quasi-orthogonal properties as DotHash. Then, Eq. (5) becomes:

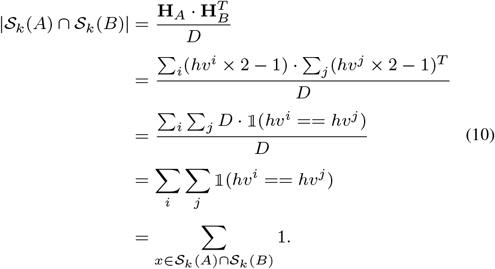

With Eq. (9) and Eq. (10), HyperGen first estimates the following Jaccard similarity using the derived sketch HVs:

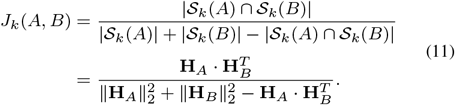

Then ANI in Eq. (8) can be easily calculated.

### 2.3 Software Implementation and Optimization

HyperGen is developed using the Rust language, and the code is available at https://github.com/wh-xu/Hyper-Gen. We present the following optimizations to improve the speed and efficiency of HyperGen.

#### 2.3.1 Sketch Quantization and Compression

Although the sketch HV has a compact data format with high memory efficiency, there still exists data redundancy in sketch HVs that can be utilized for further sketch compression. Our experimental observation is that the value range of sketch HVs is distributed within a bell curve (see Supplementary Fig. 2). Rather than store the full-precision sketch hypervector (*e*.*g*., INT32), we perform lossless compression by quantizing the HV to a lower bit width. The quantized bits are concatenated together using bit-packing.

#### 2.3.2 Fast HV Aggregation using SIMD

The inner loop of binary HV aggregation step in Algorithm 1 incurs significant runtime overhead when a large HV dimension *D* is applied. We develop a parallelized HV aggregation using single instruction, multiple data (SIMD) instruction to reduce the impact of increased HV aggregation time. As shown in Supplementary Fig. 3, the HV aggregation optimized by SIMD only takes negligible portion of the total sketching time.

#### 2.3.3 Parallel Sketching

HyperGen provides two sketching modes: 1. *normal mode* and 2. *fast mode*. The *normal mode* sketches genome files on CPU cores with multithreading. The *fast mode* offloads genome sketching to GPU with better computing capabilities. The *fast mode* can be widely supported by commodity GPUs. Our measurement results in Fig. 5 show that HyperGen’s *fast mode* further improves the sketching speed by 1.8× to 2.7× over *normal mode*.

#### 2.3.4 Pre-computation for HV Sketch Norm

The *L*^2^ norm of each sketch hypervector, ∥H∥_2_, is precomputed during sketch generation phase. The *L*^2^ norm value is stored along with the sketch hypervector to reduce redundant computations for the ANI calculation phase.

**Table 2.** Error and linearity metrics for pairwise ANI estimation. (Underline: the best among sketch-based algorithms. Bold: the best among all algorithms.)

| Dataset: *Bacillus cereus* |  |  |  |  |  |  |
| --- | --- | --- | --- | --- | --- | --- |
| Tool | $k$ | Sketch Size | MAE ↓ | RMSE ↓ | MPAE ↓ | Pearson ↑ |
| FastANI | 16 | - | **0.312** | **0.368** | **0.334** | **0.999** |
| Skani | - | 198MB (850×) | 0.354 | 0.422 | 0.377 | 0.996 |
| Mash | 21 | 830KB (3.6×) | 0.399 | 0.591 | 0.430 | 0.981 |
| Bindash | 21 | 351KB (1.5×) | <u>0.360</u> | 0.530 | <u>0.385</u> | <u>0.986</u> |
| Dashing 2 | 21 | 1.2MB (5.2×) | 0.500 | 0.650 | 0.537 | 0.981 |
| Sourmash | 21 | 11MB (47×) | 0.415 | 0.558 | 0.449 | <u>0.986</u> |
| HyperGen-2048 | 21 | 233KB (1.0×) | 0.411 | 0.707 | 0.442 | 0.975 |
| HyperGen-4096 | 21 | 459KB (2.0×) | 0.372 | <u>0.522</u> | 0.400 | <u>0.986</u> |
| Dataset: *Escherichia coli* |  |  |  |  |  |  |
| Tool | $k$ | Sketch Size | MAE ↓ | RMSE ↓ | MPAE ↓ | Pearson ↑ |
| FastANI | 16 | - | 0.680 | 1.152 | 0.705 | 0.899 |
| Skani | - | 200MB (855×) | 0.403 | 0.572 | 0.419 | **0.956** |
| Mash | 21 | 831KB (3.6×) | 0.456 | 0.686 | 0.470 | 0.930 |
| Bindash | 21 | 351KB (1.5×) | 0.442 | 0.658 | 0.456 | 0.936 |
| Dashing 2 | 21 | 1.2MB (5.1×) | 0.464 | 0.704 | 0.479 | 0.930 |
| Sourmash | 21 | 9.6MB (41×) | 0.381 | **0.565** | 0.393 | 0.944 |
| HyperGen-2048 | 21 | 234KB (1.0×) | 0.449 | 0.644 | 0.464 | 0.942 |
| HyperGen-4096 | 21 | 460KB (2.0×) | **0.368** | **0.565** | **0.381** | <u>0.952</u> |

**Table 3.** Sketch size, error, and linearity metrics for database search. (Underline: the best among sketch-based algorithms. bold: the best among all algorithms.)

| Dataset: *Bacillus cereus* |  |  |  |  |  |  |
| --- | --- | --- | --- | --- | --- | --- |
| Tool | $k$ | Sketch Size | MAE ↓ | RMSE ↓ | MPAE ↓ | Pearson ↑ |
| FastANI | 16 | - | **0.218** | **0.296** | **0.235** | **0.999** |
| Skani | 15 | 1.0GB (714×) | 0.299 | 0.378 | 0.320 | 0.998 |
| Mash | 21 | 4.7MB (3.4×) | 0.542 | 0.678 | 0.586 | <u>0.996</u> |
| Bindash | 21 | 2.0MB (1.4×) | 0.467 | 0.579 | 0.502 | 0.994 |
| Dashing 2 | 21 | 6.7MB (4.8×) | 0.576 | 0.715 | 0.622 | 0.993 |
| HyperGen-2048 | 21 | 1.4MB (1.0×) | 0.355 | 0.480 | 0.382 | 0.994 |
| HyperGen-4096 | 21 | 2.6MB (1.9×) | <u>0.318</u> | <u>0.424</u> | <u>0.342</u> | <u>0.996</u> |
| Dataset: *Escherichia coli* |  |  |  |  |  |  |
| Tool | $k$ | Sketch Size | MAE ↓ | RMSE ↓ | MPAE ↓ | Pearson ↑ |
| FastANI | 16 | - | 0.215 | 0.391 | 0.221 | 0.950 |
| Skani | 15 | 6.9GB (697×) | 0.198 | **0.277** | 0.203 | **0.983** |
| Mash | 21 | 36MB (3.6×) | 0.226 | 0.529 | 0.231 | 0.877 |
| Bindash | 21 | 16MB (1.6×) | 0.206 | 0.514 | 0.210 | 0.870 |
| Dashing 2 | 21 | 51MB (2.6×) | 0.234 | 0.536 | 0.239 | 0.873 |
| HyperGen-2048 | 21 | 9.9MB (1.0×) | 0.178 | 0.502 | 0.182 | 0.833 |
| HyperGen-4096 | 21 | 20MB (2.0×) | **0.153** | 0.491 | **0.156** | 0.851 |
| Dataset: NCBI RefSeq |  |  |  |  |  |  |
| Tool | $k$ | Sketch Size | MAE ↓ | RMSE ↓ | MPAE ↓ | Pearson ↑ |
| FastANI | 16 | - | 0.443 | 0.522 | 0.452 | 0.968 |
| Skani | 15 | 1.8GB (486×) | 0.266 | 0.292 | 0.272 | **0.997** |
| Mash | 21 | 14MB (3.8×) | 0.204 | 0.251 | 0.208 | 0.983 |
| Bindash | 21 | **5.9MB** (1.6×) | 0.238 | 0.269 | 0.243 | 0.988 |
| Dashing 2 | 21 | 20MB (5.4×) | 0.167 | 0.189 | 0.171 | 0.972 |
| HyperGen-2048 | 21 | 3.7MB (1.0×) | 0.216 | 0.304 | 0.234 | <u>0.991</u> |
| HyperGen-4096 | 21 | 7.4MB (2.0×) | **0.135** | **0.164** | **0.138** | <u>0.991</u> |
| Dataset: Parks MAGs |  |  |  |  |  |  |
| Tool | $k$ | Sketch Size | MAE ↓ | RMSE ↓ | MPAE ↓ | Pearson ↑ |
| FastANI | 16 | - | 0.457 | 0.551 | 0.490 | **0.998** |
| Skani | 15 | 6.6GB (367×) | **0.310** | **0.456** | **0.335** | 0.997 |
| Mash | 21 | 65MB (1.9×) | <u>1.090</u> | <u>1.298</u> | <u>1.137</u> | 0.990 |
| Bindash | 21 | 29MB (1.6×) | 1.096 | 1.308 | 1.140 | 0.991 |
| Dashing 2 | 21 | 93MB (5.2×) | 2.163 | 2.466 | 2.251 | 0.921 |
| HyperGen-2048 | 21 | 18MB (1.0×) | 1.291 | 1.448 | 1.374 | 0.975 |
| HyperGen-4096 | 21 | 35MB (1.9×) | 1.146 | 1.297 | 1.211 | 0.983 |
| Dataset: GTDB MAGs |  |  |  |  |  |  |
| Tool | $k$ | Sketch Size | MAE ↓ | RMSE ↓ | MPAE ↓ | Pearson ↑ |
| FastANI | 16 | - | **0.436** | **0.592** | **0.469** | 0.976 |
| Skani | 15 | 66GB (458×) | 0.466 | 0.630 | 0.500 | 0.969 |
| Mash | 21 | 533MB (3.7×) | <u>0.584</u> | <u>0.668</u> | <u>0.632</u> | **0.980** |
| Bindash | 21 | 231MB (1.6×) | 0.772 | 0.837 | 0.835 | 0.971 |
| Dashing 2 | 21 | 770MB (5.3×) | 0.994 | 1.283 | 1.078 | 0.892 |
| HyperGen-2048 | 21 | 144MB (1.0×) | 1.098 | 1.409 | 1.250 | 0.904 |
| HyperGen-4096 | 21 | 287MB (2.0×) | 0.982 | 1.138 | 1.094 | 0.974 |

## 3 Evaluation and Results

### 3.1 Evaluation Methodology

#### 3.1.1 Genome Dataset and Hardware Setting

The evaluation is conducted on a machine with a 16-core Intel i7-11700K CPU with up to 5.0GHz frequency, 2TB NVMe PCIe 4.0 storage, and 64GB of DDR4 memory. Unless otherwise specified, all programs are allowed to use 16 threads with their default parameters. Five genome datasets in Supplementary Table 2 are adopted for benchmarking. The datasets include: *Bacillus cereus, Escherichia coli*, NCBI RefSeq (Jain *et al*., 2018), Parks MAGs (Parks *et al*., 2017), and GTDB MAGs (Parks *et al*., 2018). These datasets vary in terms of number of genomes, lengths, and sizes.

#### 3.1.2 Benchmarking Tools

We compare HyperGen with five state-of-the-art tools, including Mash (Ondov *et al*., 2016), Bindash (Zhao, 2019), Sourmash (Brown and Irber, 2016), Dashing 2 (Baker and Langmead, 2023), FastANI (Jain *et al*., 2018), Skani (Shaw and Yu, 2023), and ANIm (Kurtz *et al*., 2004). Mash, Bindash, Sourmash, and Dashing 2 are sketch-based tools for ANI estimation. In comparison, FastANI and Skani use mapping-based methods while ANIm adopts the most accurate base-level alignment-based method to calculate the ANIs. ANIm results are regarded as the ground truth. Specifically, we use NUCleotide MUMmer (Kurtz *et al*., 2004) to generate the alignment results and then convert the alignment data into the corresponding ground-truth ANIs. Dashing 2 uses its weighted *bagminhash* mode. HyperGen (similar to Mash, Bindash, Sourmash, and Dashing 2) is an ANI approximation tool for the high ANI regime. We follow the previous work (Ondov *et al*., 2016) and only preserve ANI values *>* 85. The versions and commands used are summarized in Supplementary Table 1. HyperGen uses *k*-mer size *k* = 21, scaled factor *S* = 1500 as suggested in previous works (Shaw and Yu, 2023; Hera *et al*., 2023; Brown and Irber, 2016). Our analysis in Section 3.2.1 shows that the HV dimension *D* = 4096 achieves a good balance between ANI estimation error and sketching complexity. So we set it as the default parameter. HyperGen also supports the *fast mode* which accelerates the sketching process on GPU.

#### 3.1.3 Evaluation Metrics

##### ANI Precision

One of the critical metrics for evaluating the effectiveness of a genome sketching tool is the precision of ANI estimation. We use three metrics to evaluate the ANI approximation errors: 1. mean absolute error (MAE), 2. root mean squared error (RMSE), and 3. mean percentage absolute error (MPAE). We also adopt the Pearson correlation coefficient to assess the linearity of the ANI estimate with respect to ground truth.

##### Computation and Memory Efficiency

An ideal genome sketching scheme should be able to generate compact sketch files at the cost of short runtime, especially for large-scale genomic analysis. To compare the computation and memory efficiency of evaluated tools, we measure and report the wall-clock runtime and sketch sizes during database search.

### 3.2 ANI Estimation Quality

In this section, we study the quality of ANI estimation by performing the following pairwise ANI experiment. First, the largest 100 genome files are collected from each dataset. Then, each batch of 100 genome files is used to calculate the pairwise and symmetric 100 × 100 ANI matrix.

#### 3.2.1 HyperGen ANI Quality using Different Parameters

We first evaluate the impact of HyperGen’s two algorithmic parameters: scaled factor *S* and HV dimension *D* on the final ANI estimation errors and linearity. The experimental results are depicted in Fig. 3, where the scaled factor *S* and the HV dimension *D* vary from 800 to 2000 and from 256 to 16384, respectively. It shows that: for all scaled factors, the ANI approximation errors decrease significantly as *D* increases from 256 to 4096. This is because a larger HV dimension can produce better orthogonality, which is helpful to reduce the approximation error of the set intersection according to the theory in (Nunes *et al*., 2023). But increasing the HV dimension larger than *D* = 4096 does not yield a significant error reduction or linearity improvement.

**Fig. 3:**
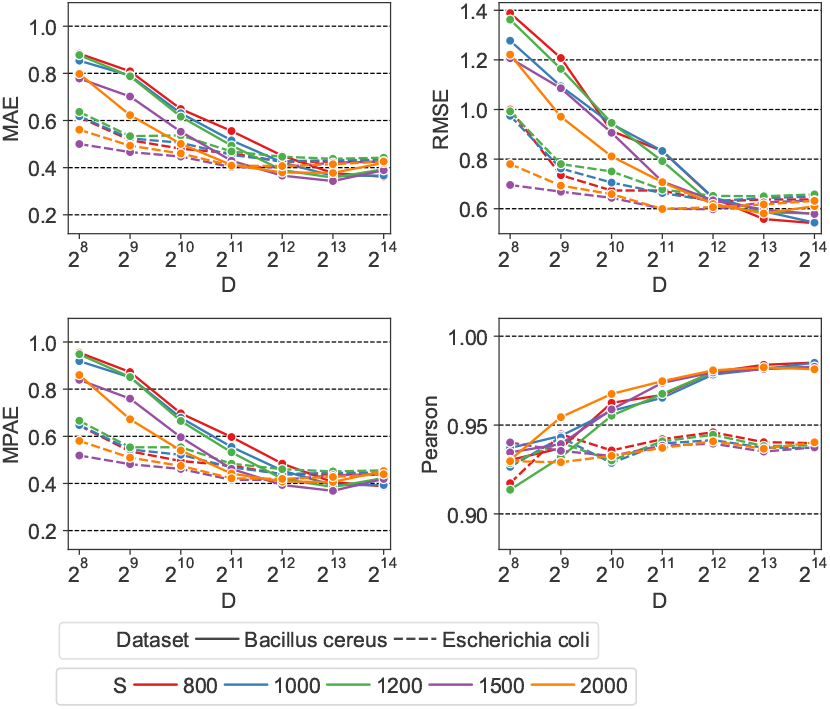
Error metrics (MAE, RMSE, MPAE) and ANI linearity (Pearson coefficient) as a function of scaled factor *S* and HV dimension *D*.

It is also observed that a smaller scaled factor *S* generally leads to a worse ANI approximation error when using the same HV dimension *D*. The reason behind this is: a smaller *S* that produces a larger hash threshold value as in Eq. (2), will generate a denser sampling of *k*-mers. This increases the size of sampled *k*-mer hash set. As a result, more binary HVs need to be aggregated to the sketch HV. The excessive number of binary HVs degrades the orthogonality between binary HVs, reducing the approximation accuracy for set cardinality. To balance between the quality and complexity of the ANI approximation, we choose *S* = 1500 and *D* = 4096 as the default scaled factor and HV dimension, respectively.

#### 3.2.2 Comparison with Other Sketching Tools

We also compare the quality of the ANI estimation for various tools, including Mash, Bindash, Dashing 2, Sourmash, FastANI and Skani. For fair comparison, the sketch-based tools (HyperGen, Mash, Bindash, Sourmash, and Dashing 2) use the same sketch size. Other parameters are the same as their default parameters. Specifically, HyperGen uses *D* = 4096, while Mash and Dashing 2 use a sketch size of 1024.

HyperGen can be used to estimate the Jaccard index. First we perform Jaccard estimation experiment and compare HyperGen to Mash, Bindash, Dashing 2, and Sourmash. Supplementary Table 3 shows the error metrics with respect to the true Jaccard results. The 100 × 100 Jaccard matrix for *Bacillus cereus* and *Escherichia coli* datasets is computed. HyperGen achieves competitive Jaccard estimation accuracy with other baseline tools.

Table 2 summarizes the ANI error and linearity metrics with respect to the ground truth values on *Bacillus cereus* and *Escherichia coli* datasets. For the *Bacillus cereus* dataset, HyperGen is slightly inferior to Bindash, FastANI and Skani, which yields a comparable Pearson correlation coefficient compared to the other sketch-based tools (Mash and Dashing 2). In the *Escherichia coli* dataset, HyperGen consistently surpasses all other sketch-based tools, providing both lower ANI approximation errors and better linearity. Meanwhile, HyperGen’s sketch size is over 800× smaller than Skani. These experiments demonstrate that HyperGen is capable of delivering a high quality of ANI estimation.

### 3.3 Genome Database Search

One critical workload that genome sketching tools can accelerate is the genome database search. Meanwhile, the genome database search can be extended to multiple downstream applications.

#### 3.3.1 ANI Linearity and Quality

We extensively consider the five evaluated datasets as reference databases. We run FastANI, Skani, Mash, Bindash, Dashing 2, and HyperGen using the commands and queries listed in Supplementary Table 2. Sourmash is not considered because it does not support multi-thread execution. The execution consists of two steps: 1. All tools first generate reference sketches for the target database, 2. The second step is to search for the query genomes against the built reference sketches. Note that FastANI and Skani were unable to complete the database search on the Parks MAGs and GTDB datasets in one shot because it requires more memory than the available 64GB and experienced out of memory issues. We divided FastANI and Skani executions into smaller batches and measured the accumulative runtime.

The estimated ANI values generated in Table 3 by each tool in the NCBI RefSeq, Parks MAGs, and GTDB MAGs datasets are depicted in Fig. 4 with their corresponding ground truth values from ANIm. Data points with ANI*<* 85 are filtered. It shows that HyperGen produces good ANI linearity compared to the ground truth results.

**Fig. 4:**
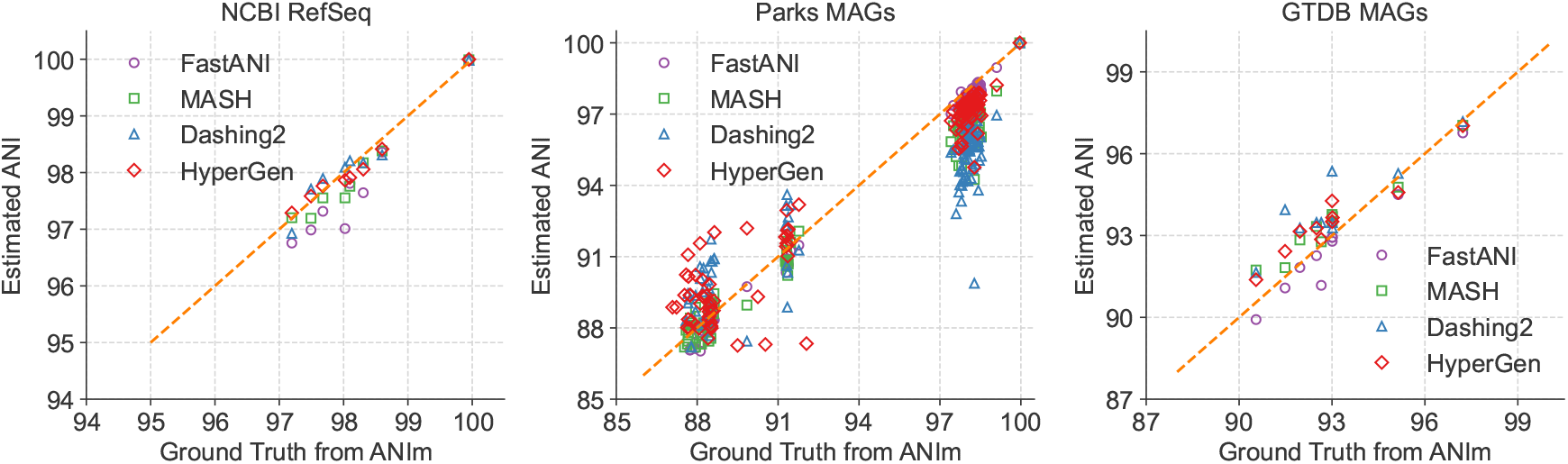
Database search ANI comparison for FastANI, Mash, Dashing 2, HyperGen, and ground-truth ANIm on NCBI RefSeq, Parks MAGs, and GTDB MAGs datasets.

Quantitative results in terms of numerical error and linearity metrics are summarized in Table 3. The ANI error distribution for each tool can be seen in Supplementary Fig. 4. In datasets *Bacillus cereus, Escherichia coli*, and NCBI RefSeq, HyperGen achieves the lowest ANI errors among all sketch-based tools, delivering more accurate ANI estimations as compared to Mash, Bindash, and Dashing 2. HyperGen still shows competitive accuracy over mapping-based FastANI and Skani. In *Escherichia coli* and NCBI RefSeq, HyperGen outperforms FastANI and Skani in terms of most error metrics and produces comparable Pearson coefficients. HyperGen is capable of achieving state-of-the-art error and linearity for large-scale genome search. Meanwhile, the required sketch size is two orders of magnitude smaller than Skani.

We study the impact of genome quality on the ANI estimation accuracy. We calculate the BUSCO completeness value (Simão *et al*., 2015) for each reference genome file. As shown in Supplementary Fig. 5, the more incomplete genomes of *GTDB MAGs* have higher ANI estimation error. Hence, applying HyperGen to incomplete genomes leads to more significant ANI errors.

#### 3.3.2 Runtime Performance

The wall-clock time spent on two major steps during database search: reference sketch generation and query search, is illustrated in Fig. 5. HyperGen-Fast means using the *fast sketching mode* on GPU. The reference sketching step is mainly bounded by the sketch generation process, while the search step is bounded by the sketch file loading and ANI calculation. HyperGen without *fast mode* achieves the 2nd fastest sketching speed, slightly slower than Skani. After enabling *fast mode*, HyperGen is the fastest sketching tool for most evaluated datasets. The sketching speed of HyperGen is 2.7× to 4.1× faster than Bindash. HyperGen is significantly faster (10× to 13×) than the mapping-based FastANI.

**Fig. 5:**
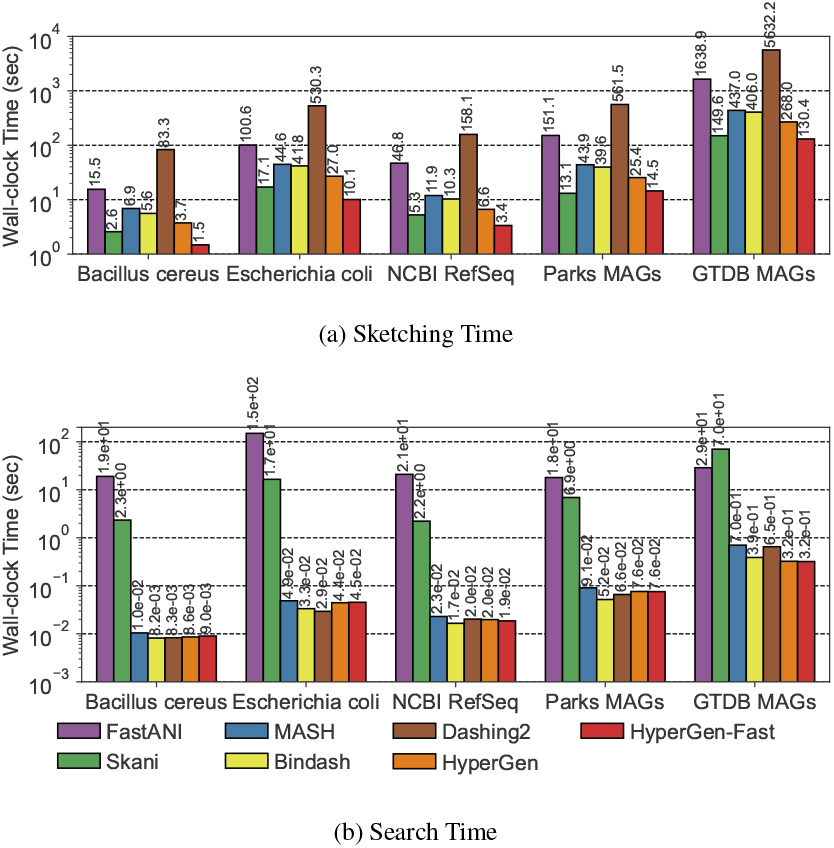
Runtime performance comparison for genome search in Table 3. (a) Reference sketching time and (b) Query search time.

For query search, HyperGen is also one of the fastest tools. The search speedup of HyperGen over FastANI and Skani is 100× to *>* 3000× because FastANI and Skani require slow sequence mapping and large index file loading processes. Moreover, the speedup of HyperGen is more significant for larger datasets. Dashing 2 sketch size is about 2.6× of HyperGen so it takes more time to load sketch files. The reduced sketch size helps to save sketch loading time. Meanwhile, the HV sketch format of HyperGen allows us to adopt highly vectorized programs to compute ANI with a short processing latency.

#### 3.3.3 Memory Efficiency

The file sizes of the reference sketches generated by Mash, Dashing 2, and HyperGen, are listed in Table 3. We apply the *Sketch Quantization and Compression* technique to HyperGen. As a result, HyperGen consumes the smallest memory space among the three sketch-based tools. The sketch sizes produced by Mash and Dashing 2 are 1.8× to 2.6× of HyperGen’s sketch sizes. This suggests that HyperGen is the most space-efficient sketching algorithm. Compared to original datasets with GB sizes, a compression ratio of 600 − 1200× can be achieved by only processing the sketch files. This enables the large-scale genome search on portable devices with memory constraints. HyperGen’s memory efficiency comes from two factors. First, the *Hyperdimensional Encoding for k-mer Hash* step converts discrete hash values into continuous high-dimensional sketch HVs, which are more compact than hash values. Second, HyperGen’s *Sketch Quantization and Compression* provides additional 1.3× compression through further removing redundant information in sketch HVs.

Table 4 summarizes performance metrics in terms of peak memory consumption and runtime for the GTDB MAG dataset search. HyperGen achieve both the fastest sketching and search speed due to the efficient HDC algorithm as well as software optimizations. FastANI and Skani experience OOM (out of memory) issues because they require a large memory space to store intermediate data for sequence mapping. In comparison, HyperGen consumes about 1GB of memory for the sketching or searching phase, significantly lower than FastANI and Skani. This indicates that HyperGen is friendly to run on memory-limited device, such as laptop.

**Table 4.** Benchmarking peak memory consumption and runtime for single-query search on GTDB MAGs dataset. OOM: out of memory.

| Tool | Sketch Phase |  | Search Phase |  |
| --- | --- | --- | --- | --- |
|  | Peak Memory | Runtime | Peak Memory | Runtime |
| FastANI | OOM | 1,638.9s | OOM | 28.8s |
| Skani | 5.3GB | 149.6s | OOM | 70.2s |
| Mash | 1.9GB | 437.0s | 1.0GB | 0.7s |
| Bindash | 0.3GB | 406.0s | 0.2GB | 0.4s |
| Dashing 2 | 8.9GB | 5,632.2s | 0.6GB | 0.7s |
| HyperGen | 1.0GB | 130.4s | 0.9GB | 0.3s |

## 4 Discussion and Conclusion

Fast and accurate estimation of Average Nucleotide Identity (ANI) is considered crucial in genomic analysis because ANI is widely adopted as a standardized measure of genome file similarity. In this work, we present HyperGen: a genome sketching tool based on hyperdimensional computing (HDC) (Nunes *et al*., 2023; Kanerva, 2009) that improves accuracy, runtime performance, and memory efficiency for large-scale genomic analysis. HyperGen inherits the advantages of both FracMinHash-based sketching (Hera *et al*., 2023; Irber *et al*., 2022) and DotHash (Nunes *et al*., 2023). HyperGen first samples the *k*-mer set using FracMinHash. Then, the discrete *k*-mer hash set is encoded into the corresponding sketch HV in hyperdimensional space. This allows the genome sketch to be presented in compact vectors without sacrificing accuracy. HyperGen software implemented in Rust language deploys vectorized routines for both sketch and search steps. The evaluation results show that HyperGen offers superior ANI estimation quality over state-of-the-art sketch-based tools (Ondov *et al*., 2016; Baker and Langmead, 2023). Meanwhile, HyperGen delivers not only the fastest sketch and search speed, but also the highest memory efficiency in terms of the sketch file size.

### Future directions of HyperGen include the following aspects

#### Further Compression and Faster Large-scale Search

The vector representation of sketch HVs allows us to apply more optimizations on the top of HyperGen. For instance, we can employ lossy vector compression techniques, such as product quantization (Jegou *et al*., 2010; Guo *et al*., 2020) and residual quantization (Lee *et al*., 2022), to reduce sketch size and memory footprint. This is advantageous for achieving rapid genome database search on embedded or mobile devices.

On the other hand, the search step in HyperGen requires intensive GEMM operations to obtain ANI values between genomes. The large-scale database search can be further accelerated using advanced hardware architectures with high data parallelism and optimized interfaces. Previous work (Xu *et al*., 2023) demonstrates that deploying HDC-based bioinformatics analysis on GPU exhibits at least one order of magnitude speedup over CPU.

#### More genome workloads

HyperGen can be extended to support a wider range of genomic applications. For example, in metagenome analysis, we can utilize HyperGen to perform the containment analysis for genome files such as (Ondov *et al*., 2019). To realize this, the sketch HVs generated by HyperGen can be used to calculate the max-containment index instead of ANI. The ANI estimation error and memory requirements of HyperGen can be reduced by considering the more accurate ANI estimation based on multi-resolution *k*-mers (Liu and Koslicki, 2022).

## Supplementary data

Supplementary data are available at Bioinformatics online.

## Data availability

The source code of HyperGen used in this work is freely available at https://github.com/wh-xu/Hyper-Gen. The scripts to reproduce the experimental results in this work can be accessed at https://github.com/wh-xu/experiment-hyper-gen. All used datasets can be downloaded from https://gtdb.ecogenomic.org and http://enve-omics.ce.gatech.edu/data/fastani.

## Footnotes

1 https://github.com/wangyi-fudan/wyhash

