## Supplementary materials for "HyperGen: Compact and Efficient Genome Sketching using Hyperdimensional Vectors"

*Bioinformatics*
doi.10.1093/bioinformatics/xxxxxx
Advance Access Publication Date: Day Month Year
Manuscript Category

Supplementary Materials

### HyperGen: Compact and Efficient Genome Sketching using Hyperdimensional Vectors

**Weihong Xu[1,*], Po-Kai Hsu[2], Niema Moshiri[1], Shimeng Yu[2], and Tajana Rosing[1]**

[1]Department of Computer Science and Engineering, University of California San Diego, La Jolla, CA 92093, USA.
[2]School of Electrical and Computer Engineering, Georgia Institute of Technology, Atlanta, GA 30332, USA.

*To whom correspondence should be addressed.

#### Abstract

This document summarizes the supplementary materials.

#### List of Supplementary Materials

- **Supplementary Table** 1
  Names, versions, and commands of benchmarked genome tools for ANI calculation. The sketch-based tools include: Mash, Dashing 2, and HyperGen. The mapping-based tool is FastANI. The alignment-based tool is ANIm.
- **Supplementary Table** 2
  Specifications (name, description, data size, query genome, and sources) of evaluated datasets.
- **Supplementary Table** 3
  Error metrics for the $100 \times 100$ pairwise Jaccard estimation. HyperGen-2048 and HyperGen-4096 use $D = 2048$ and $D = 4096$, respectively. Other tools use their default parameters.
- **Supplementary Figure** 1
  The illustration of HV orthogonality in HyperGen for HV dimension $D = 64$ to 8192 and number of elements $n = 32$ to 256.
- **Supplementary Figure** 2
  The value distribution of sketch hypervectors (HVs) generated by HyperGen when using various scaled factor $S = 800$ to 2000.
- **Supplementary Figure** 3
  The execution time breakdown of HyperGen during genome sketching. The HV dimension ranges from $D = 1024$ to 8192. The HV aggregation optimized by Single Instruction Multiple Data (SIMD) incurs negligible overhead as compared to the FracMinHash step.
- **Supplementary Figure** 4
  The ANI estimation error distribution of database search for all benchmarking tools (HyperGen, Mash, Bindash, Dashing 2, FastANI, and Skani).
- **Supplementary Figure** 5
  The relationship between completeness and absolute ANI estimation error for database search using HyperGen with parameters $k = 21, D = 4096, S = 1500$. The completeness is calculated by BUSCO (https://busco.ezlab.org/). HyperGen achieves smaller ANI estimation error for more complete genomes.

© The Author 2024. Published by Oxford University Press. All rights reserved. For permissions, please

Table 1. Names, versions, and commands of benchmarked genome tools for ANI calculation. The sketch-based tools include: Mash, Dashing 2, and HyperGen. The mapping-based tool is FastANI. The alignment-based tool is ANIm.

| Tool | Version | Commands and arguments |
|---|---|---|
| HyperGen | v0.2.2 | hyper-gen sketch -D cpu -t 16 -k 21 -s 1500 -d 4096 -p (fna_path) -o (file_out)<br>hyper-gen sketch -D gpu -t 16 -k 21 -s 1500 -d 4096 -p (fna_path) -o (file_out)<br>hyper-gen dist -t 16 -r (ref_sketch) -q (query_sketch) -o (dist_file) |
| Mash | v2.3 | mash sketch (data set) -o (sketches) -p 16<br>mash dist (query genome) (sketches) -p 16 |
| Bindash | v1.0 | bindash sketch –nthreads=16 –listfname=(genome_list) –outfname=(sketch)<br>bindash dist (query_sketch) (ref_sketch) –nthreads=16 –outfname=(dist_out) |
| Sourmash | v4.5 | sourmash sketch dna –output-dir (sketches) (data set)<br>sourmash compare (sketches)/*.sig -k 21 –max-containment –ani |
| Dashing 2 | v2.1.19 | dashing2 sketch –bagminhash -k 21 -S (sketch_size) -p 16 -F (file_list)<br>dashing2 sketch –bagminhash –cache -k 21 -S (sketch_size) -p 16 -F (ref_file) -Q (query_file) |
| Skani | v0.2.1 | skani sketch -t 16 -c 70 -m 1000 -l (genome_list) -o (sketches)<br>skani dist -t 16 -q (query_sketches) -r (ref_sketches) -o (file_out) |
| FastANI | v1.33 | fastANI –rl (genome_list) -q (query_genome) -t 16 |
| ANIm (nucmer) | v0.2.12 | average_nucleotide_identity.py -m ANIm –workers 16 -i (genomes) -o (output_folder) |

Table 2. Detailed specifications for the evaluated genome datasets.

| Dataset Name | Description | Size | Query Genome | Source |
|---|---|---|---|---|
| *Bacillus cereus* | Draft genome assemblies of *Bacillus cereus s.l.* from the prokaryote section of the NCBI Genome database. | 3.1GB | *Bacillus anthracis* (NZ_CM002395) | Dataset 2 at http://enve-omics.ce.gatech.edu/data/fastani |
| *Escherichia coli* | Draft genome assemblies of *Escherichia coli* from the prokaryote section of the NCBI Genome database. | 22GB | *Escherichia coli* (GCA_000303255) | Dataset 3 at http://enve-omics.ce.gatech.edu/data/fastani |
| NCBI RefSeq | Prokaryotic genomes downloaded from RefSeq database. | 5.6GB | *Escherichia coli* K12 W3110 (NC_007779) | Dataset 1 at http://enve-omics.ce.gatech.edu/data/fastani |
| Parks MAGs | A large collection of metagenome-assembled genomes. | 20GB | Pseudomonas stutzeri (Parks GCA_002292085_1) | Dataset 5 at http://enve-omics.ce.gatech.edu/data/fastani |
| GTDB MAGs | A phylogenetically consistent and rank normalized genome-based taxonomy for prokaryotic genomes sourced from the NCBI Assembly database. | 203GB | *Escherichia coli* K12 W3110 (NC_007779) | Release r207 at https://gtdb.ecogenomic.org/ |

Table 3. Error metrics for the 100 $\times$ 100 pairwise Jaccard estimation. HyperGen-2048 and HyperGen-4096 use $D = 2048$ and $D = 4096$, respectively. Other tools use their default parameters.

**Dataset: *Bacillus cereus***

| Tool | MAE ↓ | RMSE ↓ | MPAE ↓ |
|---|---|---|---|
| Mash | 0.008 | 0.011 | 3.860 |
| Bindash | 0.007 | 0.010 | 3.748 |
| Dashing 2 | 0.011 | 0.016 | 6.458 |
| Sourmash | 0.004 | 0.005 | 3.123 |
| HyperGen-2048 | 0.010 | 0.013 | 5.815 |
| HyperGen-4096 | 0.007 | 0.009 | 4.274 |

**Dataset: *Escherichia coli***

| Tool | MAE ↓ | RMSE ↓ | MPAE ↓ |
|---|---|---|---|
| Mash | 0.009 | 0.011 | 3.500 |
| Bindash | 0.007 | 0.009 | 2.498 |
| Dashing 2 | 0.032 | 0.051 | 8.709 |
| Sourmash | 0.013 | 0.014 | 4.394 |
| HyperGen-2048 | 0.010 | 0.013 | 3.267 |
| HyperGen-4096 | 0.009 | 0.011 | 2.641 |

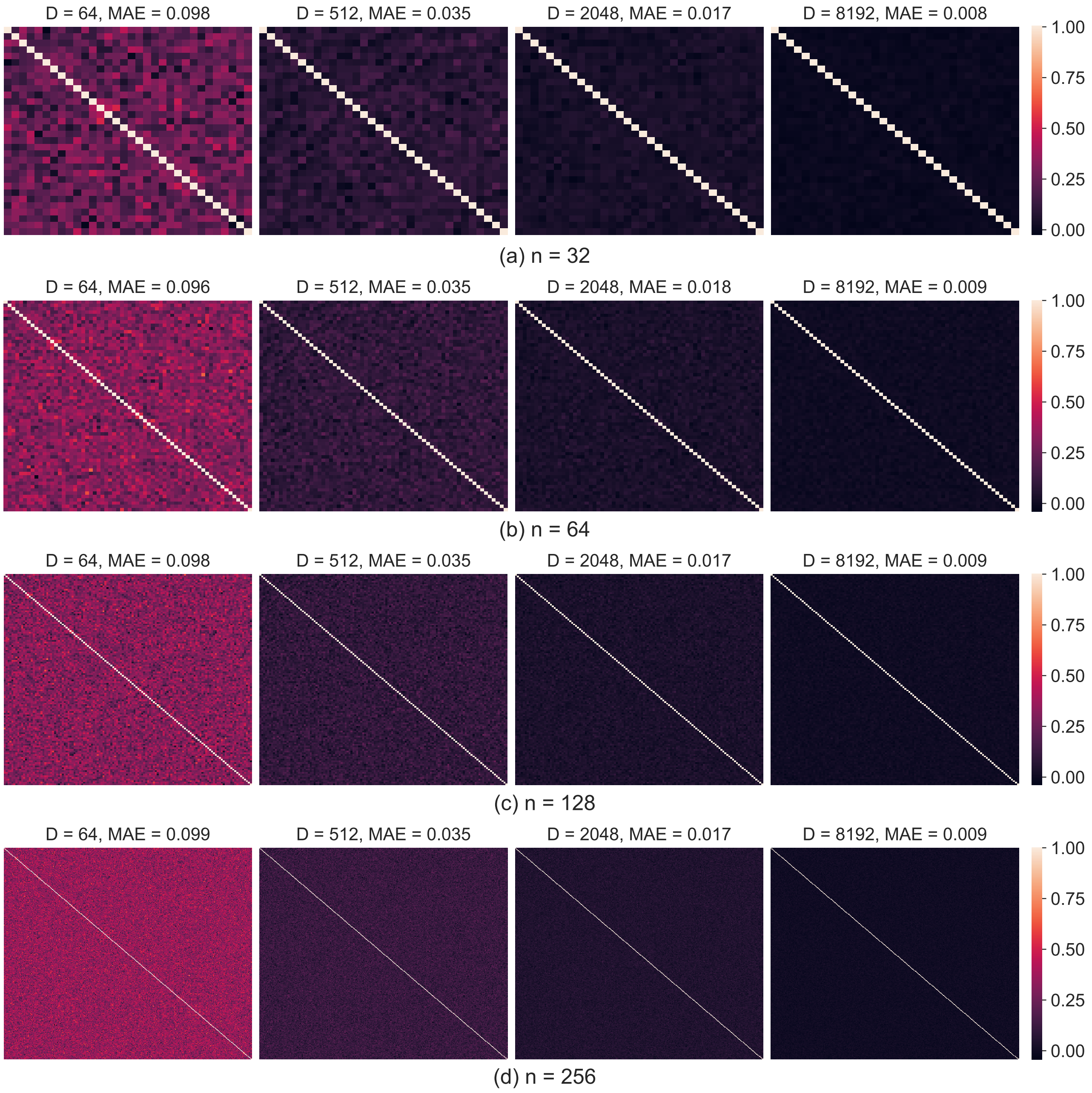


Fig. 1: The illustration of HV orthogonality in HyperGen for HV dimension $D = 64$ to 8192 and number of elements $n = 32$ to 256. The pairwise similarity for each HV is computed and depicted. HVs with the same index has similarity close to 1 while HVs with different indices are quasi-orthogonal (similarity close to 0). The mean absolute error (MAE) between the pairwise matrix and identity matrix is computed to measure the quasi-orthogonality. Larger HV dimensions provide better orthogonality (smaller MAEs).

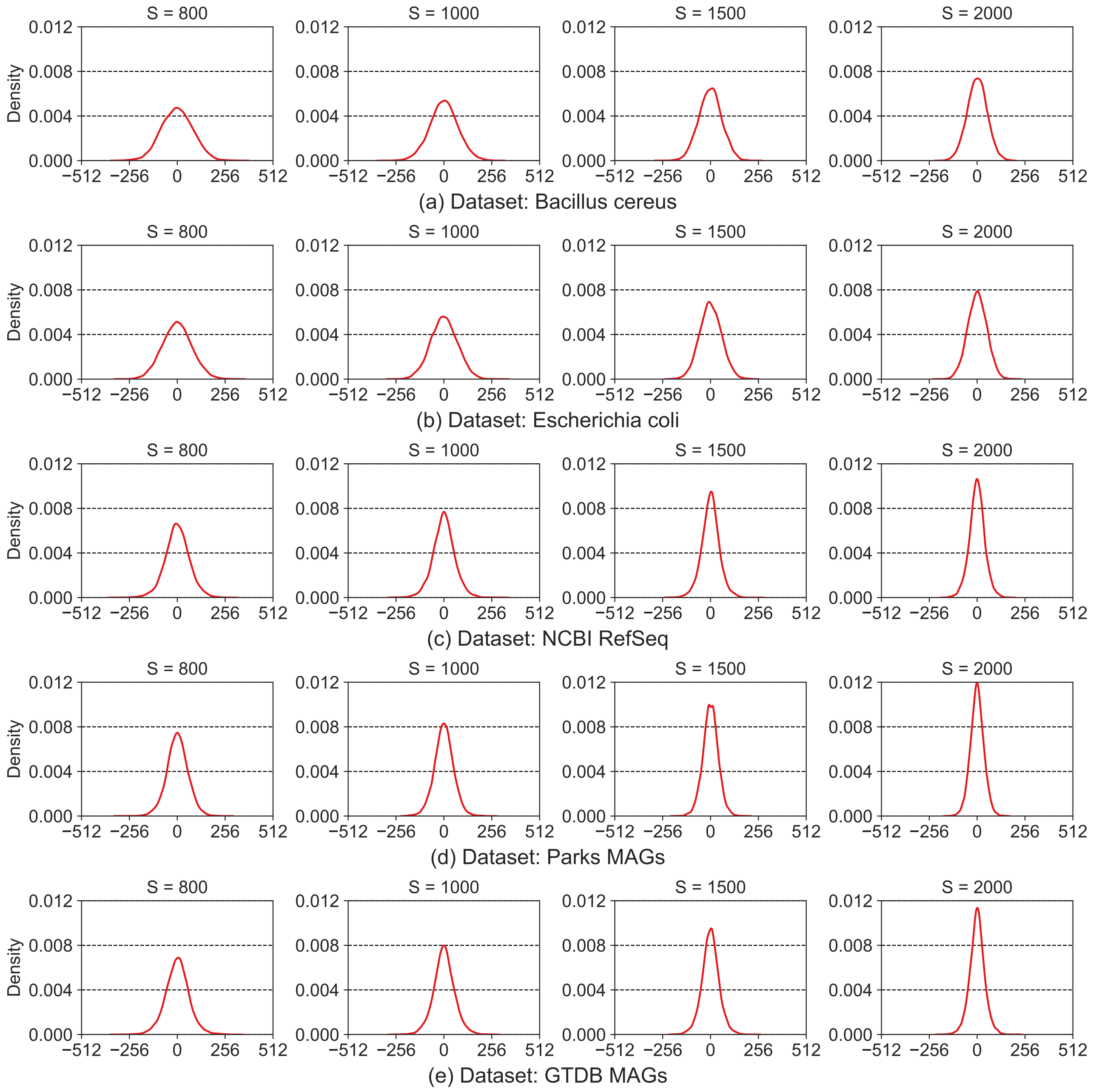


Fig. 2: The value distribution of sketch hypervectors (HVs) generated by HyperGen when using various scaled factor $S = 800$ to 2000. HV values exhibit a bell curve distribution, where the majority of values locate within the range $-300$ to 300. Sketch HVs can be effectively quantized to about 10 bits without precision loss.

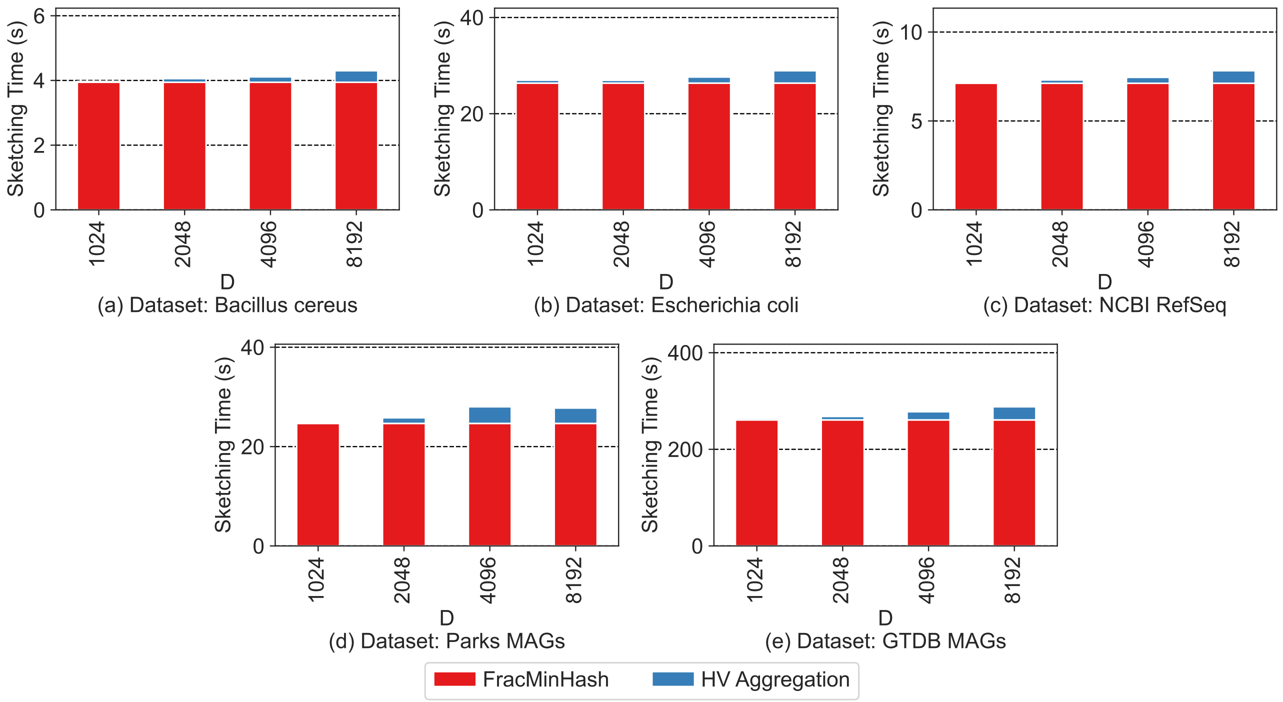


Fig. 3: The execution time breakdown of HyperGen during genome sketching. The HV dimension ranges from $D = 1024$ to 8192. The HV aggregation optimized by Single Instruction Multiple Data (SIMD) incurs negligible overhead as compared to the FracMinHash step.

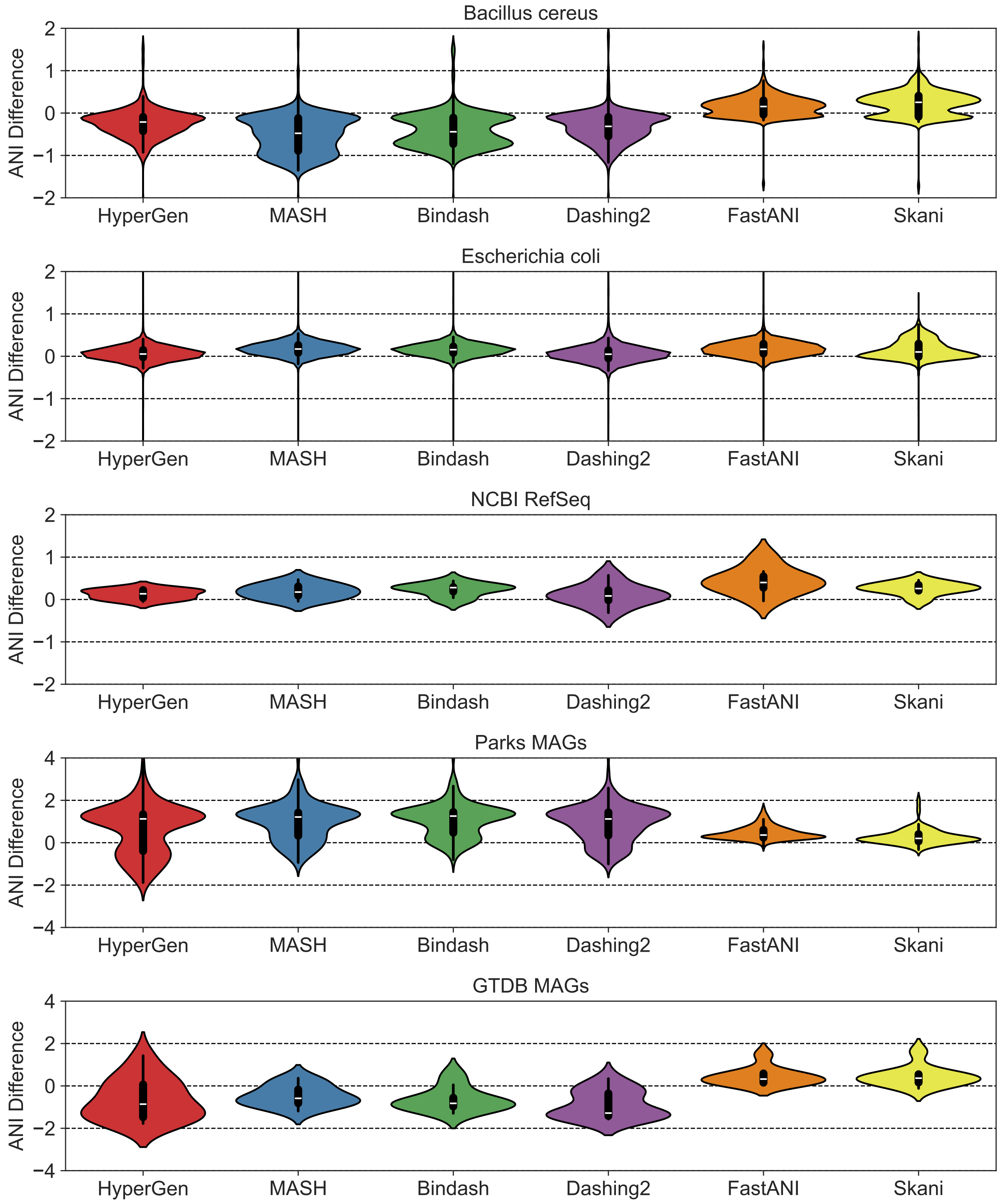


Fig. 4: The ANI estimation error distribution of database search for all benchmarking tools (HyperGen, Mash, Bindash, Dashing 2, FastANI, and Skani). Data points with ANI $> 85$ are considered here.

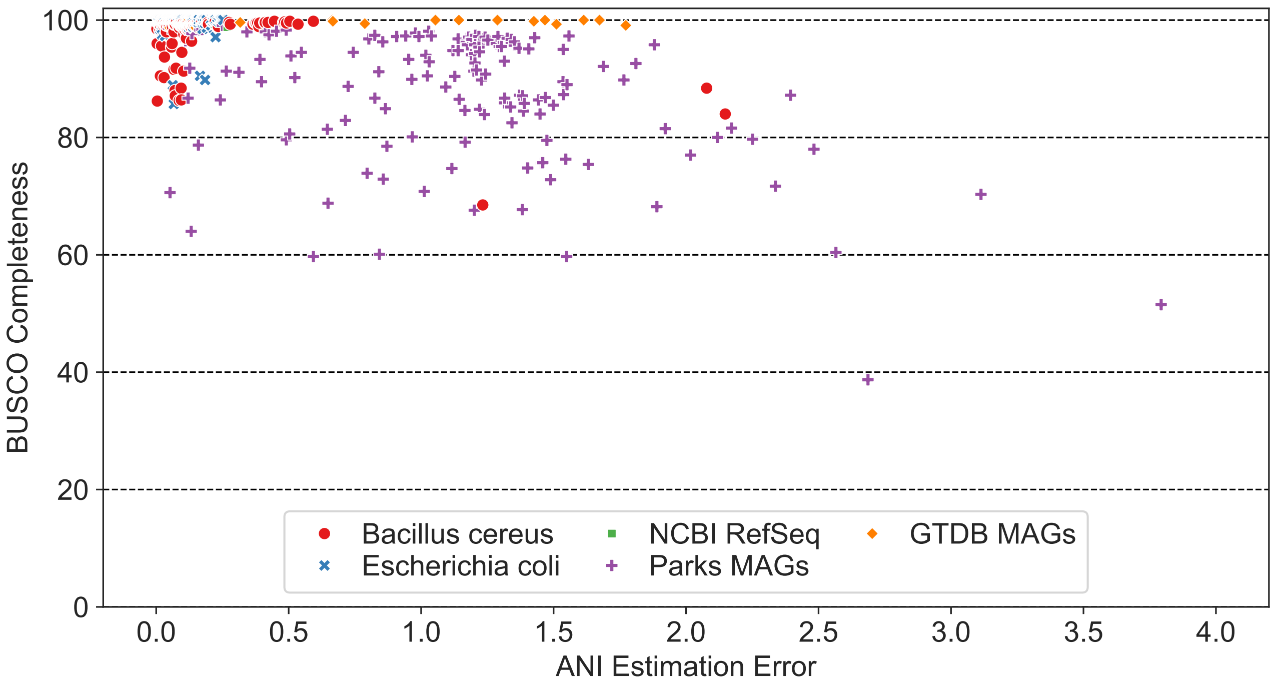


Fig. 5: The relationship between completeness and absolute ANI estimation error for database search using HyperGen with parameters $k = 21$, $D = 4096$, $S = 1500$. The completeness is calculated by BUSCO (https://busco.ezlab.org/). HyperGen achieves smaller ANI estimation error for more complete genomes. Data points with ANI $> 85$ are considered here.
